# Linking arterial biomechanics, contractility, and microstructure: A novel platform for combined structure–function assessment in murine arteries under physiological conditions

**DOI:** 10.64898/2025.12.11.692024

**Authors:** Koen W.F. van der Laan, Margarita G. Pencheva, Paul J.M. Spronck, Cédric H.G. Neutel, Pieter-Jan Guns, Casper G. Schalkwijk, Tammo Delhaas, Koen D. Reesink, Bart Spronck

**Author notes:** Corresponding author **<u>Address for correspondence</u>** Bart Spronck Department of Biomedical Engineering, Maastricht University, Universiteitssingel 40, 6229 ER, Maastricht, The Netherlands.

## Abstract

**Background:** *Ex vivo* characterization of arterial viscoelastic properties shows arterial stiffness and contractility to depend on both axial stretch and dynamic pressurization. While these arterial properties are the subject of extensive *ex vivo* research due to their relevance to vascular pathophysiology, only few experimental approaches mimic both physiological axial stretch and dynamic pressurization when characterizing arterial biomechanics, vasoreactivity, and tissue microstructure. To fill this gap, we developed a custom dynamic biaxial pressure myograph compatible with two-photon laser scanning microscopy (TPLSM).

**Methods:** We studied five murine carotid artery segments. Sample viscoelastic behaviour was characterized by quasi-static and dynamic pressurization experiments at and around physiological axial stretch, as well as quasi-static stretching at physiological pressures. In addition, vasoconstriction in response to 2 µM phenylephrine was measured during dynamic pressurization and with axial loads that mimicked physiological conditions. Lastly, arterial collagen, elastin, and cell nuclei were imaged using TPLSM with the sample at physiological axial stretch and pressurized at 100 mmHg.

**Results:** The setup enabled capture of the non-linear biaxial viscoelastic behaviour of the arterial wall as well as the viscoelastic stiffening with dynamic pressurization. Modulation of these characteristics upon stimulated smooth muscle contraction was also captured well. Moreover, the related ultrastructural properties of the collagen-elastin network as well as the transmural cell distribution, were recordable at corresponding loading conditions by TPLSM.

**Conclusion:** The presented multi-modal characterization platform enables comprehensive *ex vivo* measurements under well-controlled *in vivo*-like loading conditions, for in-depth studies focusing on arterial stiffening. Our findings emphasize the need for controlling dynamic pressure and axial stretch conditions in investigating mechanistic and constitutive aspects of arterial stiffening.

## Introduction

Arterial stiffening is a strong risk factor for cardiovascular diseases, it occurs naturally with age, and is accelerated by hypertension and diabetes (1-3). Healthy arteries are compliant at low blood pressures, stiffen at high blood pressures to prevent rupture, and regulate blood flow through contraction of the artery’s diameter (4-7). However, as arteries age, gradual degradation of elastin, accumulation of collagen, and altered contractility of vascular smooth muscle cells (VSMCs) result in stiffer arteries and disturbed arterial vasoreactivity (8-13). This stiffening reduces the damping effect of arteries on pulsatile pressure generated by the heart, which in turn can damage downstream organs (14-17).

### From assessment in vivo to understanding pathophysiology

Pulse wave velocity (PWV) is inherently related to the stiffness of the arterial wall and is the gold standard for quantifying arterial stiffening in the clinic (18). However, clinical methods for measuring PWV have little control over physiological conditions such as heart rate, blood pressure, diet, stress, etc. Because the biomechanical behaviour of arteries depends on these conditions, PWV as a metric for arterial stiffness provides limited information on the underlying changes to arterial wall constituents leading to arterial stiffening. Consequently, *ex vivo* investigations of arterial stiffness are the subject of extensive research, as they employ various types of biomechanical testers that provide greater control over experimental conditions (18-21). These setups measure arterial tissue deformations resulting from carefully applied mechanical loads under well-controlled thermal and vasoactive conditions, investigating biomechanical parameters, such as PWV and VSMC contractility, as well as biaxial stretch, stress, and stiffness.

### Relevance of physiological mechanical loading

Arterial biomechanics and VSMC contractility significantly depend on biaxial loading conditions (22-24), demonstrating the importance of mimicking physiological biaxial loading conditions when investigating arterial stiffness using *ex vivo* experiments. Furthermore, pulsatile pressurization, as compared to quasi-static pressurization, has been shown to affect arterial biomechanics and vasoreactivity through viscoelastic stiffening and VSMC contractility modulation (22, 25), respectively. While some studies report investigating arterial biomechanics or VSMC contractility while mimicking pulsatile pressurization conditions (22, 26), these conditions are generally not mimicked during *ex vivo* experiments that investigate arterial stiffening. This is predominantly due to practical restrictions, as most biomechanical testers do not have the capacity to accurately apply and control fast variations in applied circumferential load.

### Relationships between macro and microstructure and function

Like arterial biomechanics and vasoreactivity, microscopy research characterizing arterial wall microstructure benefits from mimicking physiological loading conditions, as the microstructural organization of arterial wall constituents will change under different biaxial loading conditions (11, 27). The advantages of mimicking physiological loading conditions are further emphasized when investigating correlations between changes in arterial microstructure, biomechanics, and vasoreactivity, as changes in these aspects of the arterial wall can be more directly linked to each other. However, to our knowledge, no methodology yet exists that simultaneously investigates arterial biomechanics, microstructure, and VSMC contractility while mimicking physiological biaxial and pulsatile loading conditions. Because of the lack of such integrated data, it remains unclear how *in vivo* changes in one aspect of arterial stiffening influence other aspects. Thus, interpretations around causality and vicious cycles (of causes and effects) remain cumbersome.

Therefore, the aim of the present paper is to describe and demonstrate the capabilities of our custom platform for investigating — under controlled biaxial and pulsatile loading conditions — arterial viscoelastic properties, microstructure, and VSMC contractility. We will present experimental data obtained in five mice, to showcase the potential of the platform in a research context.

## Methods

### Setup

To perform our biomechanical characterization experiments we developed a custom biaxial pressure myograph capable of quasi-static and dynamic pressurization, motorized sample stretching, and vasoreactivity experiments (**Figure 1**). The setup is globally divided into two parts, the organ bath sub-assembly that houses the sample and the surrounding control hardware (**Figure 1**), with the organ bath sub-assembly being mountable under a laser-scanning microscope.

**Figure 1:**
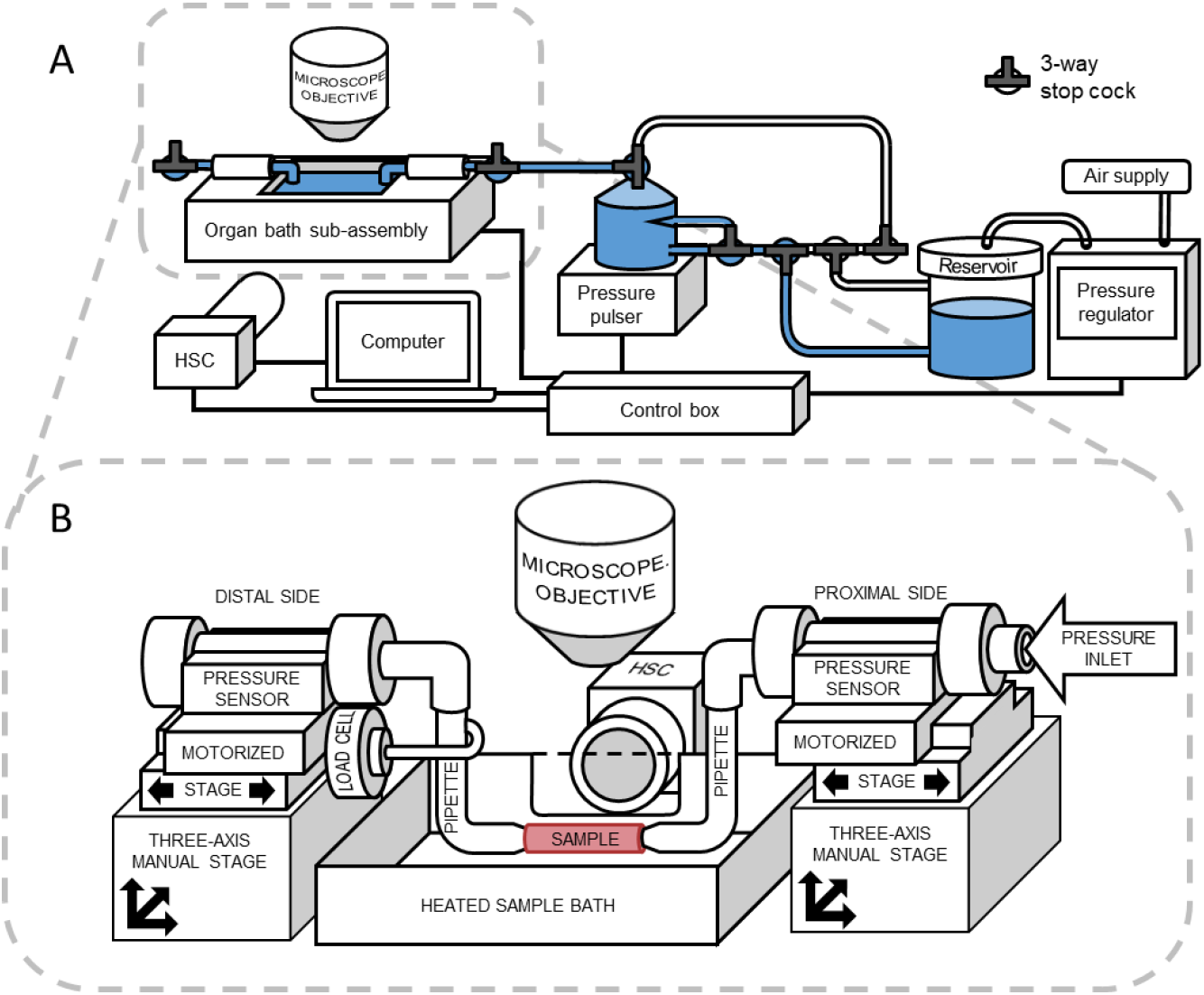
A) Biaxial biomechanical testing setup with high-speed-camera (HSC). B) Movable organ bath sub-assembly, with motorized and positionable mounting pipettes for mounting samples.

### Organ bath sub-assembly

The organ bath sub-assembly houses the sample in a heated organ bath; controls sample position and stretch; and measures luminal pressure and axial load (**Figure 1B**). Samples are mounted on L-shaped glass pipettes with roughened ends whose outer diameter slightly exceeds the inner diameter of unloaded samples, thereby preventing the samples from slipping off when loaded. The range of pipette diameters allows testing of samples ranging from mouse carotid arteries to rat thoracic descending aortas. A proximal and distal three-axis manual positioning stage aligns pipette ends so that the sample lies flat and perpendicular to the optical axis of the diameter tracking camera. A proximal and distal TRA25PPD motorized stage (Newport, Irvine, California, United States) axially stretch samples during experiments. By stretching the samples from both ends, the centre of sample remains within the field of view of the camera during stretching. In addition, a proximal and distal Transpac IV pressure transducer (Medexsupply, New Haven, Connecticut, United States) measure luminal pressures during experiments. The distal pressure sensor is immediately followed by a closed end that keeps the connection between the sensor and the sample as short and rigid as possible, with the incompressible luminal volume behind the sample ensuring pressure is measured without any time delay [{Bruggen, 2021 #243}]. A model 31 load cell (50 gram, Honeywell, Columbus, Ohio, United States) measures the axial force acting on the sample. The load cell is connected to the distal pipette with a metal hook and because the distal pipette is mounted in a flexible tube, acting as a pivot point, the pipette transmits axial forces to the load cell. Lastly, the organ bath includes a heating element and temperature sensor for controlling buffer temperature.

The connection between the organ bath sub-assembly and the rest of the setup was made flexible so that just the sub-assembly could be placed on the optical table of a confocal microscope. This allows the setup to investigate arterial wall microstructure using two-photon laser-scanning microscopy (TPLSM) as well as characterize arterial wall biomechanics. Two-photon fluorescence (TPF) and second harmonic generation (SHG) allow for imaging of specific arterial wall constituents in three dimensions while the sample is statically pressurized and axially stretched.

### Control hardware

The surrounding hardware provides static and pulsed pressure to the organ bath sub-assembly, tracks sample diameter, reads out sensors, and drives hardware (**Figure 1**). A PCD-5PSIG-D-PCV30 pressure regulator (Alicat Scientific, Tucson, Arizona, United States), connected to a buffer reservoir, provides quasi-static pressure to the setup. In addition, a custom pressure pulser generates pulsed pressure, centred on the quasi-static pressure set by the regulator. The pressure pulser generates oscillating pressure by applying force from an electrical coil against a flexible membrane that acts as the bottom of the pulser reservoir. Because the pulser generates pressure oscillations proportional to an electrical signal, the setup can generate customizable pressure oscillations. A three-way tap at the bottom of the pulser reservoir connects it to the buffer reservoir either through regular tubing or through a needle connection (**Figure 1A**). Regular tubing minimizes the resistance between the reservoirs, minimizing the pressurization response time between the sample and pressure regulator when quasi-statically pressurizing samples. In contrast, the needle connection maximizes the resistance between the two reservoirs, preventing the pressure regulator from flattening the pulsed pressure generated by the pressure pulser whilst still maintaining the average pressure. The combination of a pressure generator and pulser makes the setup capable of quasi-statically pressurizing samples from 0 to 250 mmHg and dynamically pressurizing them with a peak-to-peak pressure up to 40 mmHg with pulse frequencies up to 20 Hz, as measured by the distal pressure sensor.

The hardware control box reads out data from the pressure and temperature sensors, as well as the load cell, using a USB-6001 data acquisition card (DAQ) (National Instruments, Austin, Texas, United States) at 2000 Hz. In addition, the DAQ drives the heating pad and provides commands to the pressure regulator and pulser. The hardware control box drives the motorized stages in the sub-assembly with a TMCM-6110 motion control card (Trinamic, Eindhoven, the Netherlands). An AUI-3040CP-M-GL Rev.2 high-speed camera (HSC) (IDS Imaging Development Systems, Farnham, England) attached to an InfiniProbe TS-160 lens (Infinity Photo-Optical Company, Centennial, Colorado, United States) is used to track sample diameter through a glass window in the side of the organ bath (**Figure 1A**). The HSC takes frames with a resolution of 1448 by 542 pixels^2^ from 1 to 500 frames per second. An electrical flash trigger is provided by the HSC to the DAQ to synchronize the two data streams during data processing. Each flash trigger lasts 1 millisecond, or 2 DAQ data-points, and is generated at the start of each frame recorded by the HSC.

### Calibration

Camera zoom was preset to a fixed magnification during experiments, with the magnification depending on the used animal model, ensuring that the full sample width remains within the imaging frame. Pixel sizes for preset magnifications were calibrated by imaging an object of known size whilst ensuring that the object was located where the sample would be in the setup and that the HSC was focused on the object. Motorized stages were calibrated using a procedure provided by the motion control card that sweeps the motors between end stops. The pressure sensors were calibrated using the pressure regulator, which in turn was calibrated and certified by its manufacturer. Care was taken to account for the pressure generated by the height difference between water levels in the organ bath and buffer reservoir when calibrating the pressure sensors. Lastly, the load cell was calibrated *in situ* by connecting a spring of known stiffness to the pipettes and using the motorized stages to controllably stretch the spring.

### Experimental case protocol

Adult (5-month-old, n=5) male C57BL/6J mice (Charles River Laboratories, Lyon, France) were used for this study to demonstrate the capabilities of the setup. All animals were housed in the animal facility of the University of Antwerp in standard cages with 12h-12h light-dark cycles and had free access to regular chow and tap water. Mice were euthanized by perforating the diaphragm (performed under deep anaesthesia). Afterwards, the thoracic descending aorta was carefully removed from the mouse, placed in HEPES saline buffer, and transported to Maastricht University on ice. Samples were stripped from all adherent tissue and side branches were sutured closed. All animal experiments were approved by the Ethical Committee of the University of Antwerp (ECD n° 2022/43) and were conducted in accordance with the EU Directive 2010/63/EU.

#### Vasoconstriction

The setup and organ bath were filled with HEPES buffered saline solution (**Table 1**), taken care to remove any bubbles from the setup and keeping the organ bath at 37 C°. The sample was sutured onto the glass pipettes and brought to its unloaded length, at which the sample was neither buckled nor stretched. After checking that the sample contained no leaks and maintained luminal pressure, the sample was brought to its *in vivo*-like length. This was done by pressurizing samples with a sinusoidal pressure wave, ranging from 60 to 140 mmHg with a pulse frequency of 1 Hz, and lengthening them until the variation in measured axial force was minimal. The sample was then dynamically pressurized with a sinusoidal pressure wave ranging from 70 to 110 mmHg at a frequency of 10 Hz. Using live diameter tracking, these pressurization conditions were maintained until the sample exhibited a stable pressure–diameter cycle.

**Table 1:**
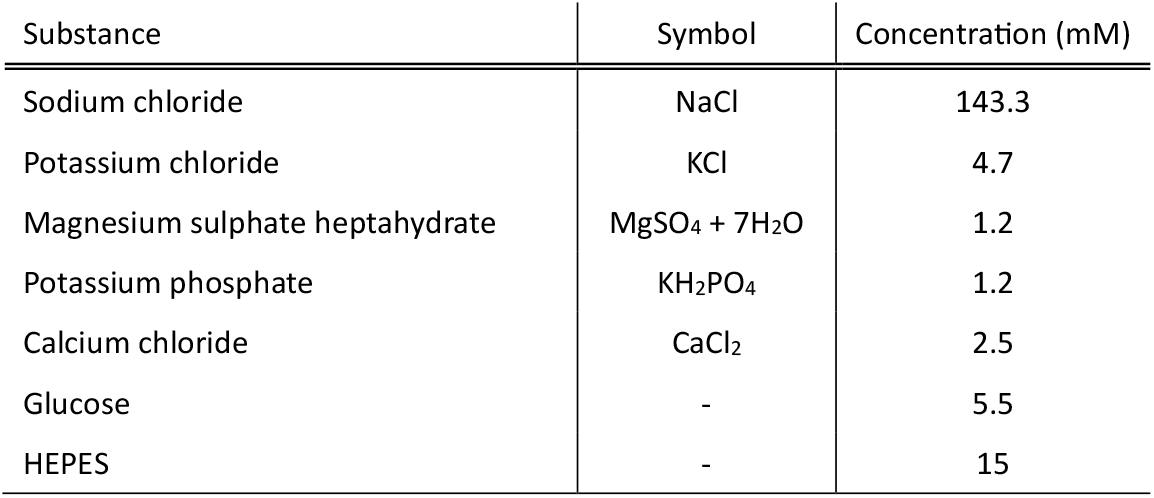
Recipe for HEPES buffered saline solution.

Contraction was induced by adding 2 µM of phenylephrine to the organ bath, after which the sample was given thirty minutes to contract. During this period, the high-speed camera recorded frames at a rate of 1.015 Hz, to not run out of memory during the experiment. Because the imaging period is not a full multiple of the pressure pulse period, each frame images a different moment of the pressure-diameter cycle. For the given pulse and frame rates, each frame advances 0.0148 seconds along the pressure-diameter cycle, equal to a phase shift of 0.930 radians, thus slowly capturing the entire cycle within seven subsequent frames. However, seven frames advance slightly more than a full period along the pressure-diameter cycle, advancing 0.1036 seconds instead of 0.1 for a full period, or 6.503 radians instead of 2*π* = 6.283 radians. The following seven frames will thus image the next pressure–diameter cycle at slightly shifted points compared to the previous cycle.

#### Biomechanical characterization

After the vasoconstriction experiment, the sample was brought to an unloaded state. The buffer in the setup was replaced with calcium- and magnesium-deficient Hanks’ buffered salt solution (HBSS) buffer at 37 C°, with the addition of 10 µM sodium nitroprusside (SNP), to suppress contraction during biomechanical characterization. Then, the sample’s unloaded length and *in vivo*-like length were re-determined. A circumferential preconditioning step followed where the sample was stretched to 105% of their newly estimated *in vivo*-like length and pressurized from 10 to 180 mmHg four times. During the following axial preconditioning step, samples were pressurized to 100 mmHg and axially stretched four times from zero force to the maximum axial force measured during circumferential preconditioning. After preconditioning, unloaded and *in vivo*-like lengths were re-determined. Passive biomechanical characterization protocol was performed in three steps: quasi-static pressurization, dynamic pressurization, and finally quasi-static axial stretching. Throughout quasi-static pressurization, samples underwent pressurization in duplicate, ranging from 10 to 180 mmHg and then back, at a rate of 3 mmHg per second. Simultaneously, they were stretched to 105, 95, and 100% of *in vivo*-like length, with the HSC set to 3 frames per second. During dynamic pressurization samples were kept at *in vivo*-like length and subjected to sinusoidal pressure waves. Samples were dynamically pressurized using the pressure pulse frequencies 0.625, 1.25, 2.5, 5, 10, and 20 Hz, for the pressure ranges of 40 to 80 mmHg, 80 to 120 mmHg, and 120 to 160 mmHg, with the HSC set to 500 frames per second. Lastly, quasi-static axial stretching experiments were performed in duplicate, stretching samples from zero force to maximum axial force measured during quasi-static pressurization experiments. Experiments involving stretching were performed at 10, 60, 100, 140, and 180 mmHg. The HSC set to 5 frames per second, and the stretching rate was set at 0.0187 s^-1^ relative to the unloaded length of the sample.

#### Two-photon microscopy imaging

To dye elastin fibres, 125 nM Eosin-y was added to the organ bath, following biomechanical characterization, and left to incubate for 20 minutes. Samples were stretched to their *in vivo*-like length, pressurized to 100 mmHg, and the removable organ bath subassembly was placed under a Radiance2100 two-photon laser-scanning microscope (TPLSM) [Bio-Rad, Hercules, California, United States]. The tuneable Tsunami pulsed laser [Spectra-Physics, Milpitas, California, United States] was set to 810 nm. 3D Image stacks were taken using a 60x, 1.00 NA objective [Nikon, Tokyo, Japan] for a field of view of 205×205 µm^2^ with a resolution of 1024×1024 pixels^2^, and a step size of 0.45 µm between images. Image stack depth was set to penetrate the sample until all image contrast was lost.

For the first image stack, second harmonic generation (SHG) and two-photon fluorescence (TPF) image modalities were used to image collagen and elastin fibres. A filter block consisting of a 500 nm dichroic mirror together with a 405-10 nm and 535-50nm fluorescent filter was used to split and direct the backscattered SHG and TPF light to two non-descanned detectors. To stain DNA, 1.5 µM of Hoechst-3342 was added to the organ bath after the first image stack and left to incubate for 60 minutes. For the second image stack, a filter block consisting of a 530 nm dichroic mirror together with a 460-50 nm and 560-40 nm fluorescent filter was used to split and direct the backscattered Hoechst-3342 and Eosin-y TPF light to two non-descanned detectors.

#### Vessel cross section imaging

After microscopy imaging, samples were depressurized, brought to their unloaded length and three thin rings were cut from the centre of the sample. Care was taken to cut the rings as perpendicular to the sample centreline as possible. These rings were then submerged in a petri dish containing calcium and magnesium deficient HBSS buffer and a millimetre measuring scale. Using a stereo microscope with adjustable zoom and an AM423X USB camera (Dino-Lite, Torrance, California, United States), cross sections of the sample were imaged (**Figure 2A**).

**Figure 2:**
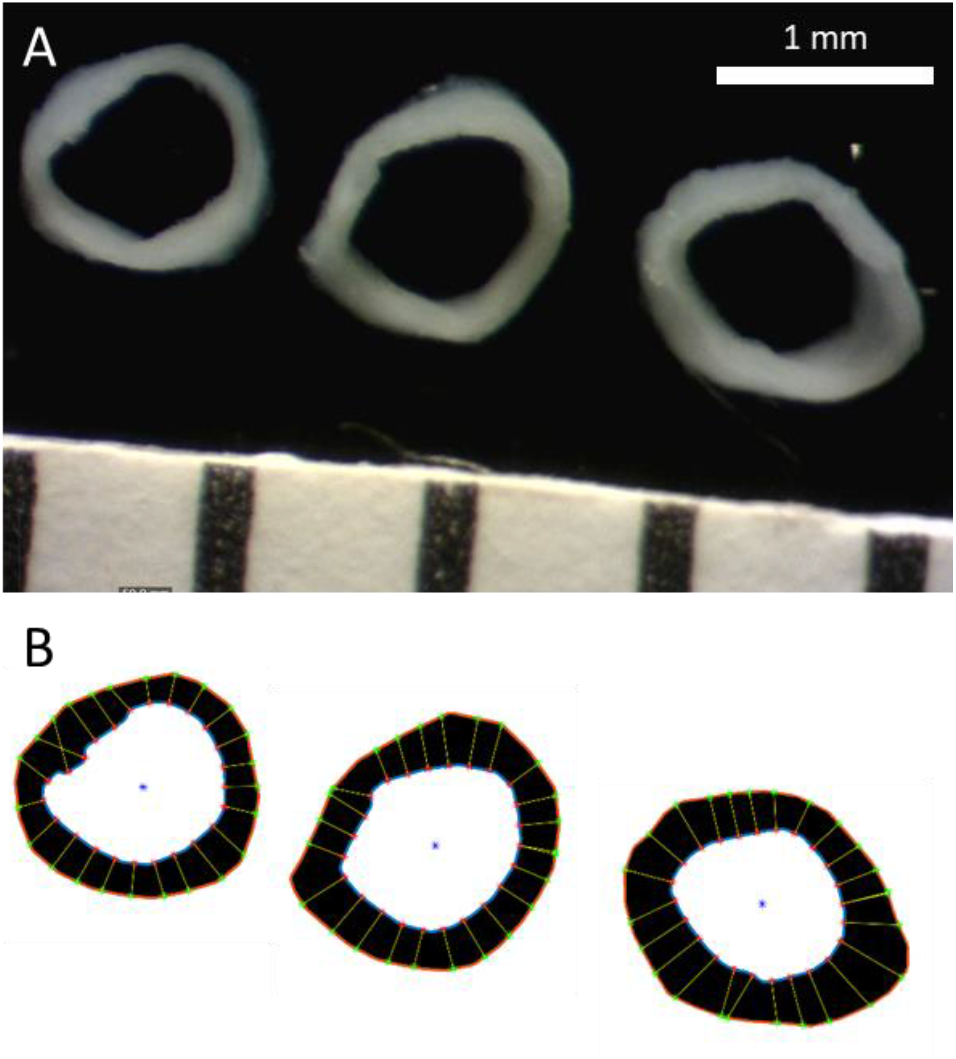
A) Cross-sectional rings cut from the sample for determining wall thickness. B) Image analysis results of determining wall thickness along 24 points per ring.

### Data processing

#### Outer diameter tracking

A custom MATLAB [MathWorks, Natick, Massachusetts, United States] image analysis script was deployed to track sample diameters from the recorded videos (**Figure 3**). First, pixel intensities were limited to a maximum of 16% of the maximum intensity (**Figure 3B**). This was to eliminate any bright spots from the video as these could be mistaken for the edge of the sample. Then, a 2-D convolution was applied to each frame using a kernel specialized in vertical edge detection (**Figure 3D**). The 40×40 pixel^2^ kernel consisted of a 2D Gaussian curve with the standard deviations along both axes set to 13 pixels and the left half of the kernel set as negative values (**Figure 3C**). The kernel was normalized so that the absolute volume beneath its surface equalled one. The maximum and minimum values along each row of the convoluted frames corresponded to the edges of the sample (**Figure 3E**). Sample diameter for the duration of the video was tracked by averaging the distance between the maximum and minimum values along each row per convoluted frame. Absolute sample diameter was then calculated by multiplying the tracked diameter in pixels with the calibrated pixel size.

**Figure 3:**
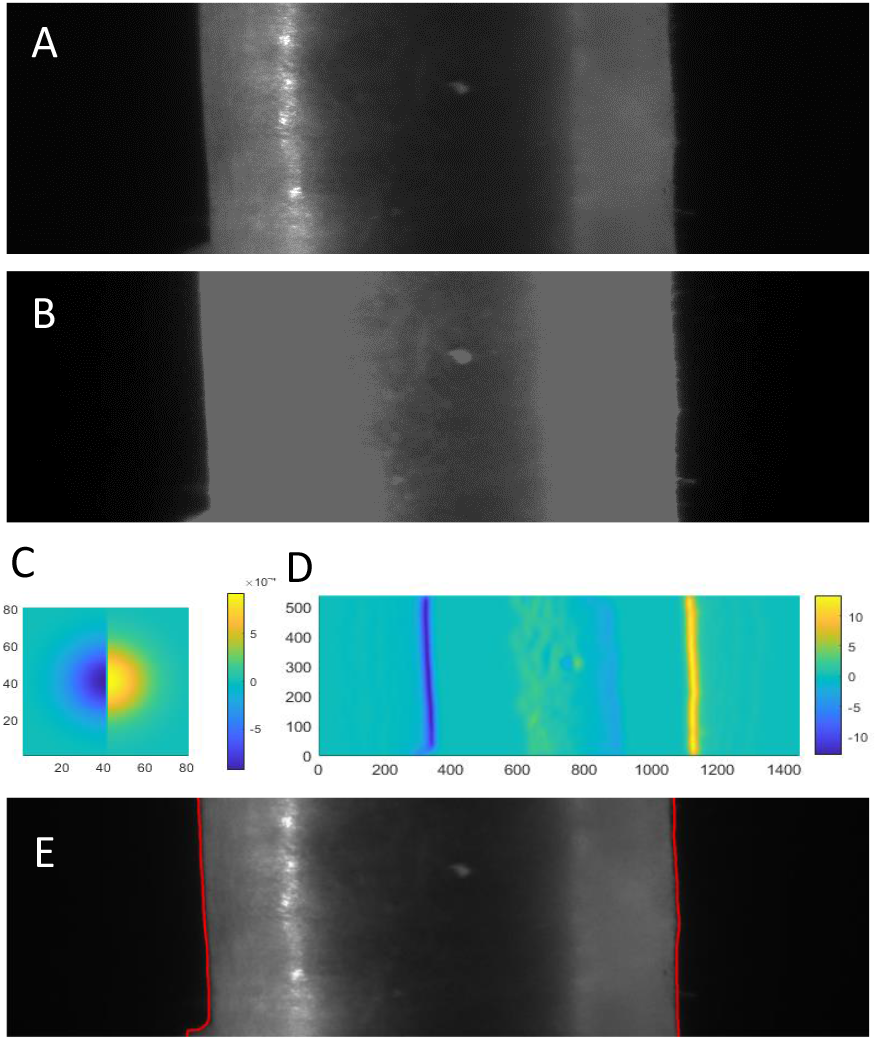
High speed camera (HSC) image processing steps for outer diameter tracking. Raw HSC frames (A) had bright spots removed using a maximum intensity threshold (B). Using a vertical edge detection kernel (C), sample edges were enhanced using 2-D convolution (D). E) Minimum and maximum values along each pixel row (shown in red) marked sample edge positions.

#### Inner diameter conversion

A custom MATLAB script was used to determine the sample wall thickness from images of cross section rings image. First, pixel size is calibrated based on the imaged millimetre scale. Then, the inner and outer edges are manually tracked for each ring. Wall thickness was determined at 24 equally spaced locations along the inner edge of each ring by the line, perpendicular to the inner edge, from the inner to outer edge (**Figure 2B**). Sample wall thickness was determined by the average of the ring wall thicknesses.

Assuming an incompressible arterial wall, conservation of volume is used to determine inner diameters from outer diameters and axial lengths measured during all vasoconstriction and biomechanical characterization experiments. Given the known dimensions of samples in their unloaded state, inner diameters were calculated according to:

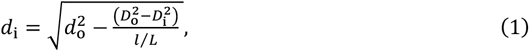

where *d*_i_ and *d*_o_ represent the loaded inner and outer sample diameters, respectively, and *l* the samples length. Unloaded sample length, inner and outer diameter are shown by *L, D*_*i*_, and *D*_*o*_, respectively.

#### Data synchronization

The flash triggers recorded by the DAQ were used to synchronize the diameter data stream with the pressure, load, and stretch data streams. The first flash trigger determines the starting time point of the diameter data, with each subsequent flash trigger determining the time point for each subsequent diameter data point. This eliminates any time delays between diameter, pressure, and axial load data, considering the timing between them is determined by the same internal clock. Stretch data was acquired through the motion control card and was synchronized to the other data streams using the computer’s internal clock. Considering stretch experiments were performed at quasi-static stretch rates, synchronization is less critical.

#### Contraction tracking

Because vasoconstriction was induced under pulsatile pressure conditions, contraction was defined as the change in inner diameter at the mean pressure during the experiment, i.e. at 90 mmHg. Due to the low frame rate, there is no guarantee a diameter was recorded at 90 mmHg. Hence, the initial diameter at 90 mmHg was determined by the intersection point of the line between the two nearest data points above and below 90 mmHg amongst the first seven data points in the experiment. The final diameter at 90 mmHg was determined in the same way but instead used the last seven data points in the experiment.

#### Biomechanical parameterization

Wall thickness, axial stress, hoop stress, and PWV were determined from quasi-static pressurization and stretch experiments to characterize arterial elastic behaviour. To minimize viscoelastic effects, present during quasi-static mechanical loading of tissue, deformation response to loading and unloading were averaged to approximate a pure elastic response. Wall thickness was determined by the difference between the sample outer and inner radius. Axial stress, *σ*_*a*_, was calculated according to the equation:

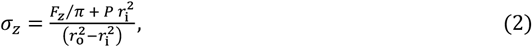

where *F*_*z*_, *P, r*_o_, and *r*_i_ represent the measured axial force, luminal pressure, outer radius, and inner radius, respectively. Circumferential stress was calculated according to the equation:

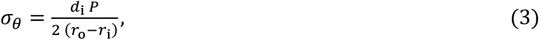

PWV was calculated using the Bramwell-Hill equation according to:

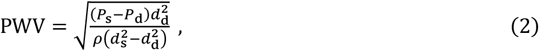

where *P*_s_ and *P*_d_ represent systolic and diastolic luminal pressures, respectively, and *d*_s_ and *d*_d_ represent systolic and diastolic inner diameter. Blood density (*ρ*) was set to 1060 kg/m^3^.

### Statistical analysis

A one-way analysis of variance (ANOVA) statistical test was used to determine the significance of the differences between PWVs determined from quasi-static and dynamic pressurization experiments. *P*-values below 0.05 were considered significant.

## Results

### Quasi-static biomechanical characterization

The combined quasi-static biomechanical characterization experiments captured the nonlinear biaxial elastic behaviour of the arterial wall (**Figure 4**), illustrating the interdependence between luminal pressure, measured axial force, inner diameter, and axial length.

**Figure 4:**
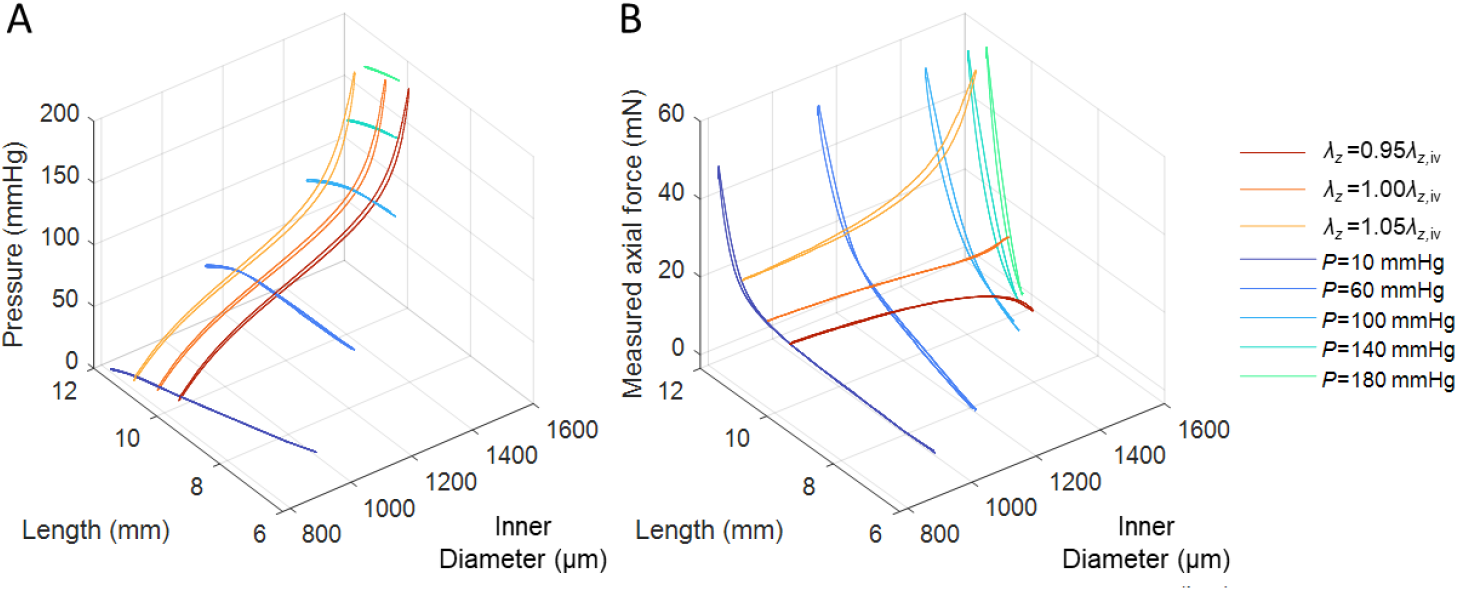
Measured aortic quasi-static biomechanics show nonlinear behaviour, due to the interdependence of axial and circumferential behaviour. A) Aortic pressure-length-inner diameter behaviour. B) Aortic measured axial force-length-inner diameter behaviour. Quasi-static inflation results are shown in yellow-to-red, while quasi-static stretch results are shown in blue-to-green.

#### Pressure-dependent biomechanical properties

Pressure-dependent biomechanical behaviour shows the non-linear elastic behaviour of arteries that progressively becomes stiffer when reaching physiological luminal pressures (**Figure 5**). This is demonstrated by the inner diameter increasing sharply with pressure at low pressures but increasing less at higher pressures (**Figure 5A**). Similarly, wall thickness decreases sharply with pressure at low luminal pressures but decreases less at higher pressures (**Figure 5B**). In contrast, PWV decreases slightly with pressures until 60 mmHg, after which the trend switches and it increases sharply with pressures (**Figure 5F**). Circumferential stress increases with pressure, slowly at first but increasingly so as pressure increases (**Figure 5E**).

**Figure 5:**
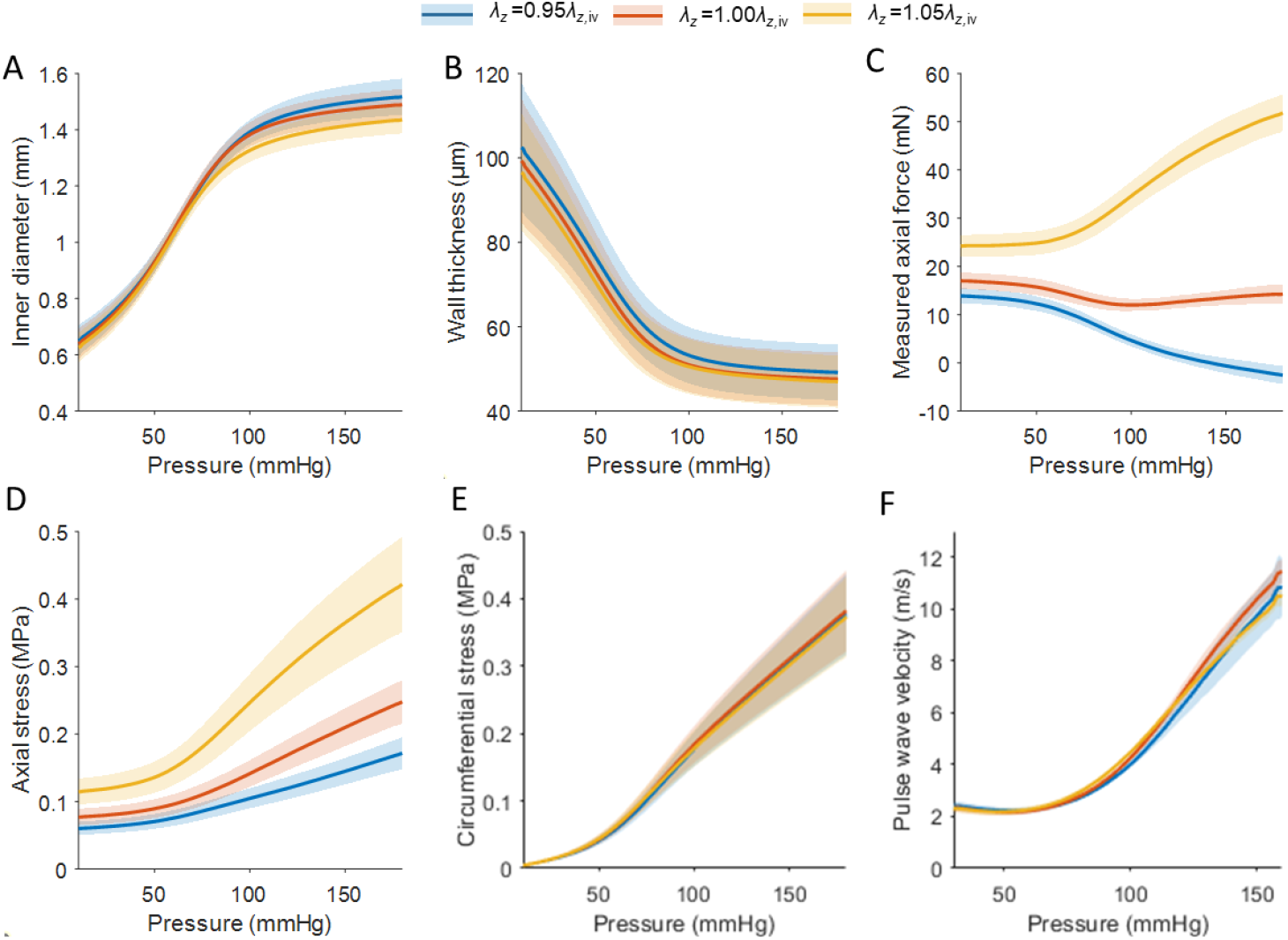
Measured biomechanical parameters demonstrate nonlinear behaviour of arterial circumferential biomechanics, stiffening as pressure reaches physiological pressures. A) inner diameter, B) wall thickness, E) circumferential stress, and F) pulse wave velocity (PWV) were largely independent of axial stretch, compared to their dependence on pressure. C) measured axial force and D) axial stress showed a strong dependence on both pressure and axial stretch.

When samples were at their *in vivo*-like axial length, measured axial force remained relatively constant, showing a small dip at 90 mmHg (**Figure 5C**). However, when samples were kept at a length shorter than their *in vivo*-like axial length, measured axial force decreases with increasing pressure. The opposite held true when samples were kept at a length longer than their *in vivo*-like axial length, with measured axial force increasing with increasing pressure. Axial stress only increased with increasing pressure, with the degree of increase dependent on axial stretch (**Figure 5D**).

#### Stretch-dependent biomechanical properties

Quasi-static stretching results demonstrate how a larger range of axial stretches affected arterial biomechanical properties, compared to quasi-static pressurization experiments (**Figure 6**). Inner diameter decreases with increasing axial stretch, except at low stretch when pressurized to 60 mmHg (**Figure 6A**). Wall thickness decreases with axial stretch until samples reach 1.06 times their *in vivo*-like length, after which wall thickness increases (**Figure 6B**).

**Figure 6:**
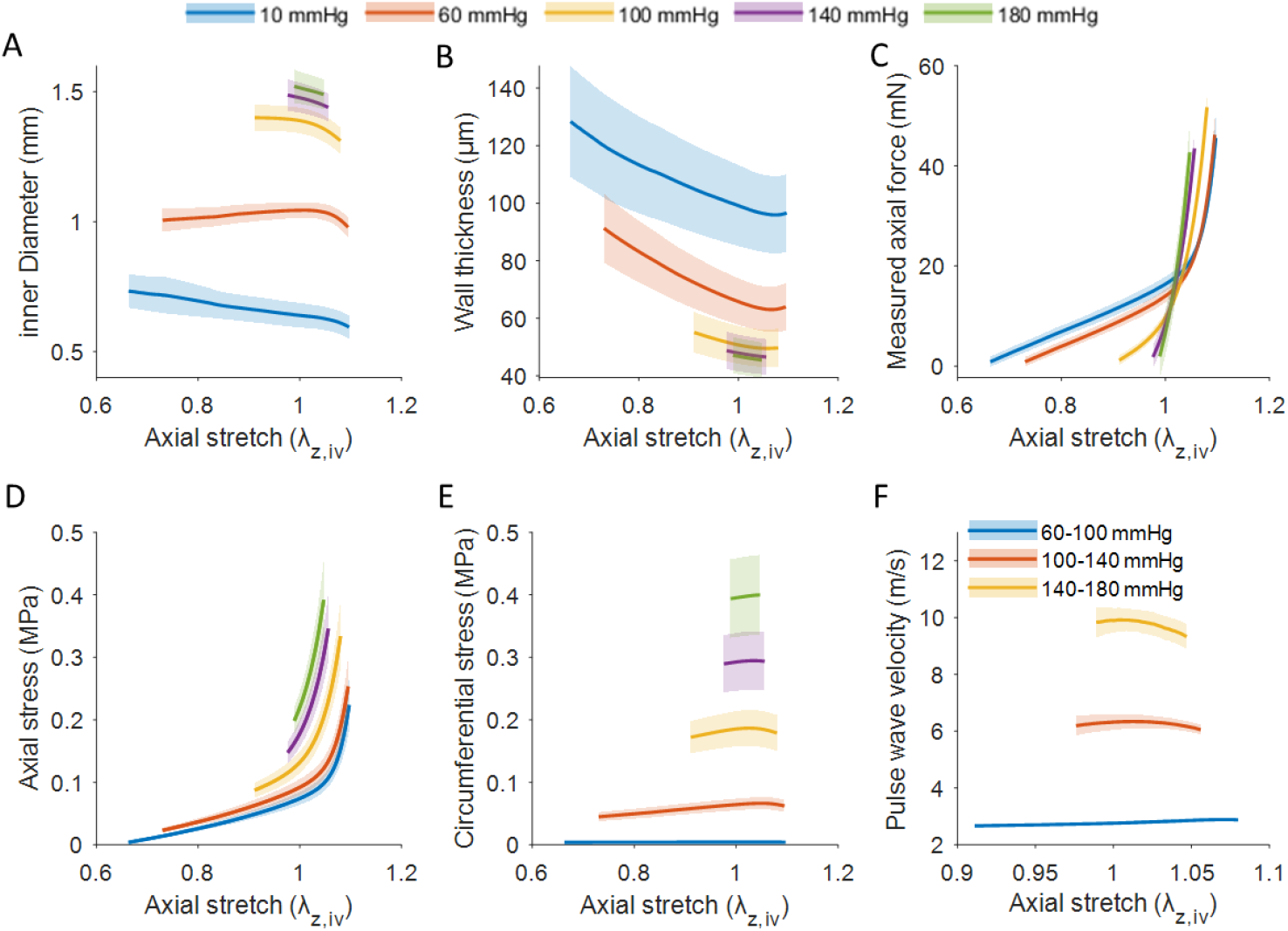
A) inner diameter and B) wall thickness decrease with axial stretch, except when samples were pressurized to 60 mmHg or exceeding a stretch of 1.06, respectively. C) Measured axial force and D) axial stress increased with axial stretch, progressively with increasing pressure. E) Circumferential stretch and F) pulse wave velocity (PWV) were largely independent of axial stretch, compared to pressurization.

Measured axial force increases with increasing axial stretch (**Figure 6C**). At low luminal pressures axial force displays a bimodal dependency on axial stretch, slowly increasing at first while sharply increasing when stretched beyond *in vivo*-like length. This behaviour transitions to a more linear relationship with increasing pressure, showing a strong dependence of measured axial force on axial stretch. The measured axial force curves intersect roughly at *in vivo*-like axial length. Axial stress shows nonlinear dependence on axial stress, increasing sharply when stretched to and beyond *in vivo*-like axial length (**Figure 6D**). The stretch at which axial stress sharply increases, decreases with increasing luminal pressure, to the point that the tipping point is no longer in the measured stretch range at 180 mmHg. As seen in quasi-static pressurization results, circumferential stress seems unaffected by axial stretch near *in vivo*-like axial stretch (**Figure 6E**). When shortening samples beyond 0.95% *in vivo*-like axial stretch, circumferential stress decreases. Similarly, PWV seems unaffected by axial stretch at low pressures but shows a slightly more curved behaviour at higher pressures, peaking at *in vivo*-like axial stretch (**Figure 6F**).

### Dynamic biomechanics

Viscoelastic stiffening causes a steepening of sample’s pressure-inner diameter behaviour when pressurizing dynamically instead of quasi-statically (**Figure 7A-C**). Consequently, dynamic pressurization experiments show that PWV increased significantly for all tested pressure pulse frequencies and pressure ranges, compared to PWV derived from quasi-static pressurization, with p< 0.001 for all pressure pulse frequencies (**Figure 7D**). In contrast, there were no significant differences between PWVs for different pressure pulse frequencies with corresponding pressure ranges.

**Figure 7:**
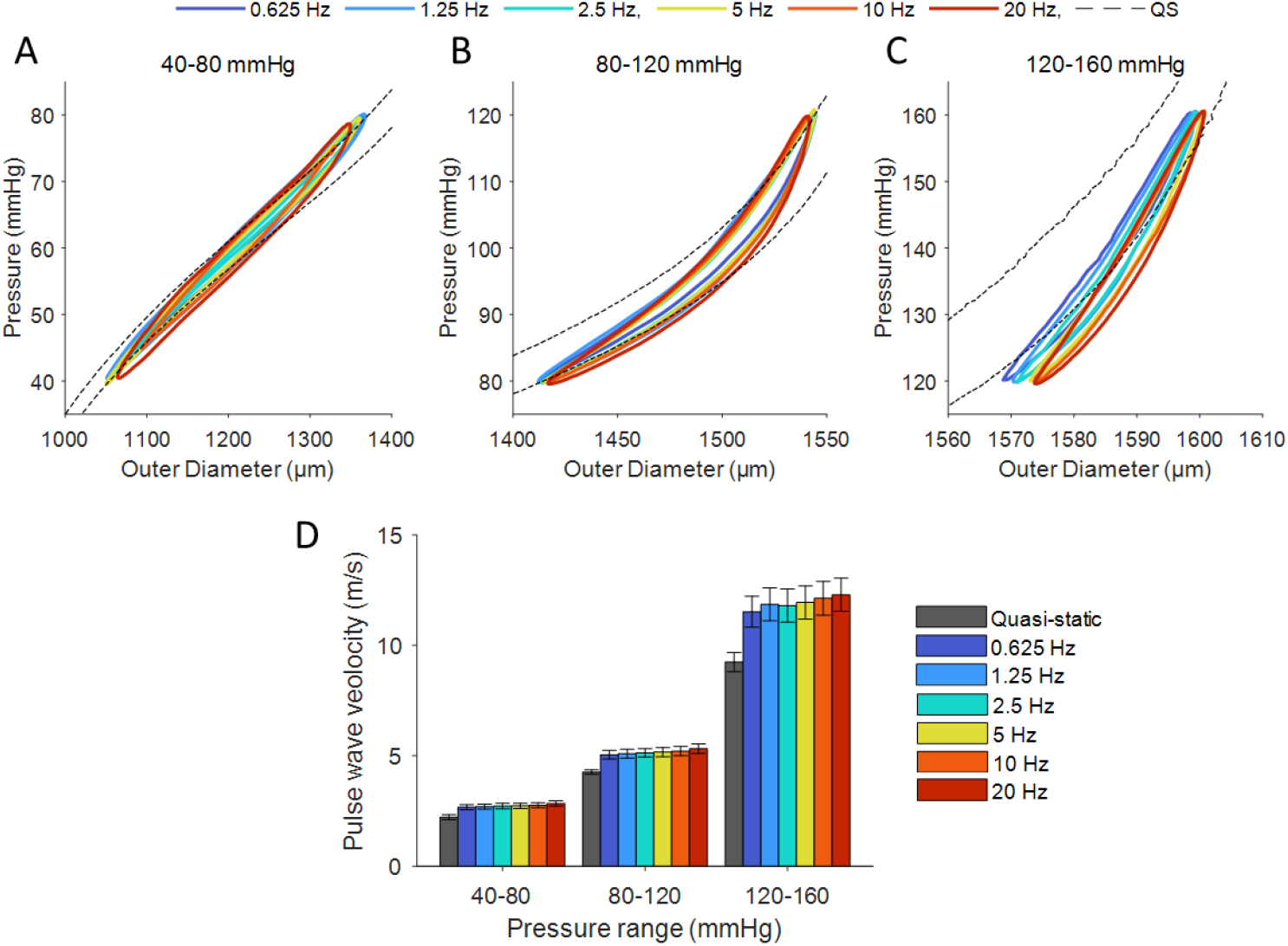
Dynamic pressure-outer diameter loops with ranges of (A) 40-80, (B) 80-120, and (C) 120-160 mmHg display steeper slopes than quasi-static pressure-outer diameter loops for the same pressure ranges. D: Pulse wave velocities are increased during dynamic pressurization, showing an increasing trend with increasing pressure pulse frequency.

### Vasoconstriction

The slight mismatch between frame time and the nearest multiple of the pressure pulse period made it possible to continuously track the dynamic pressure-inner diameter behaviour of the arterial wall during the 30-minute vasoconstriction experiment (**Figure 8A**). Samples display a clear diameter reduction in response to adding phenylephrine to the organ bath during dynamic pressurization (**Figure 8B**). Beyond a reduction in diameter, contraction inconsistently altered the observed pressure oscillation range during the vasoconstriction experiments, increasing or decreasing either the diastolic or systolic pressure, despite setup operating parameters being kept constant during the experiments.

**Figure 8:**
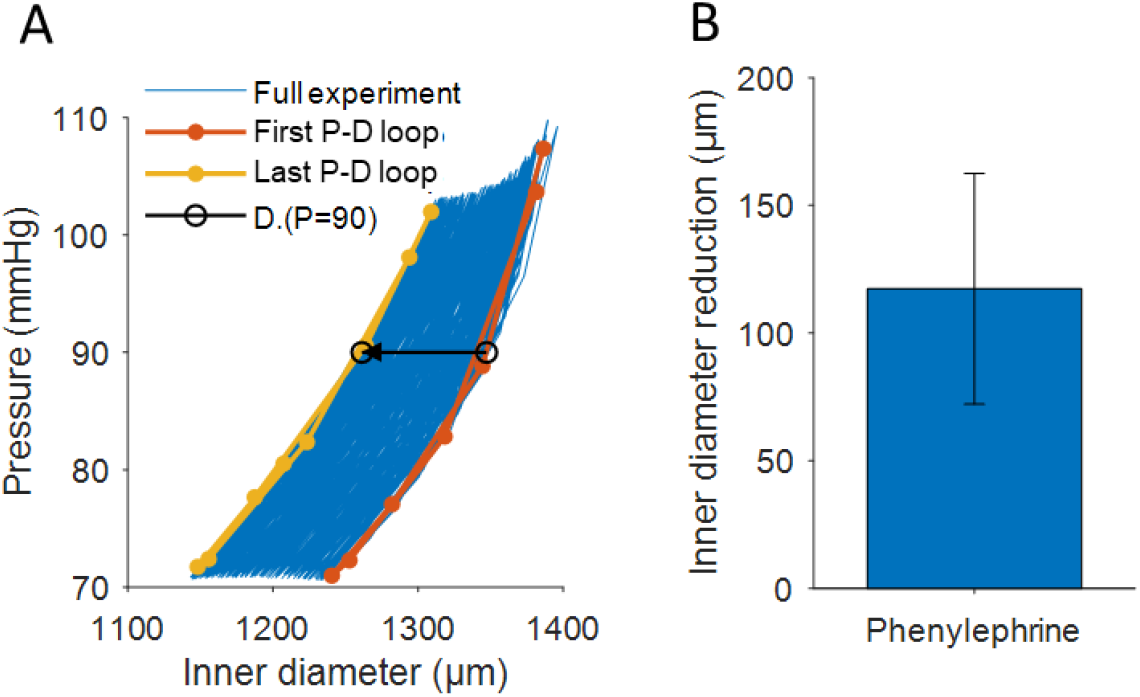
A: Sample inner diameter response to phenylephrine during vasoconstriction experiment (shown in blue), whilst dynamically pressurized. Diameter reduction at 90 mmHg (shown in black) is determined by where the first and last fully measured pressure diameter cycles (shown in orange and yellow, respectively) cross 90 mmHg. B: Inner diameter contraction response to phenylephrine during vasoconstriction experiment at 90 mmHg.

### Two-photon microscopy

TPLSM image stacks show the collagen and elastin structure within the sample wall (**Figure 9A, B**), with a thin layer of collagen fibre bundles on the outer side of the sample wall followed by a thicker layer comprised of an elastin fibre mesh. While some elastin fibres could be seen in the outer layer of collagen fibre bundles, no collagen fibres were visible in the inner elastin mesh layer. The elastin fibre mesh appeared to display a layered structure where elastin intensity alternates radially (**Figure 9B**). The second series of image stacks displayed similar elastin structures as the first (**Figure 9C, D**). In addition, Hoechst-3342 showed elongated nuclei throughout the sample wall, primarily situated between the elastin layers (**Figure 9C, D**). The nuclei laid flat inside the sample wall and were primarily circumferentially oriented.

**Figure 9:**
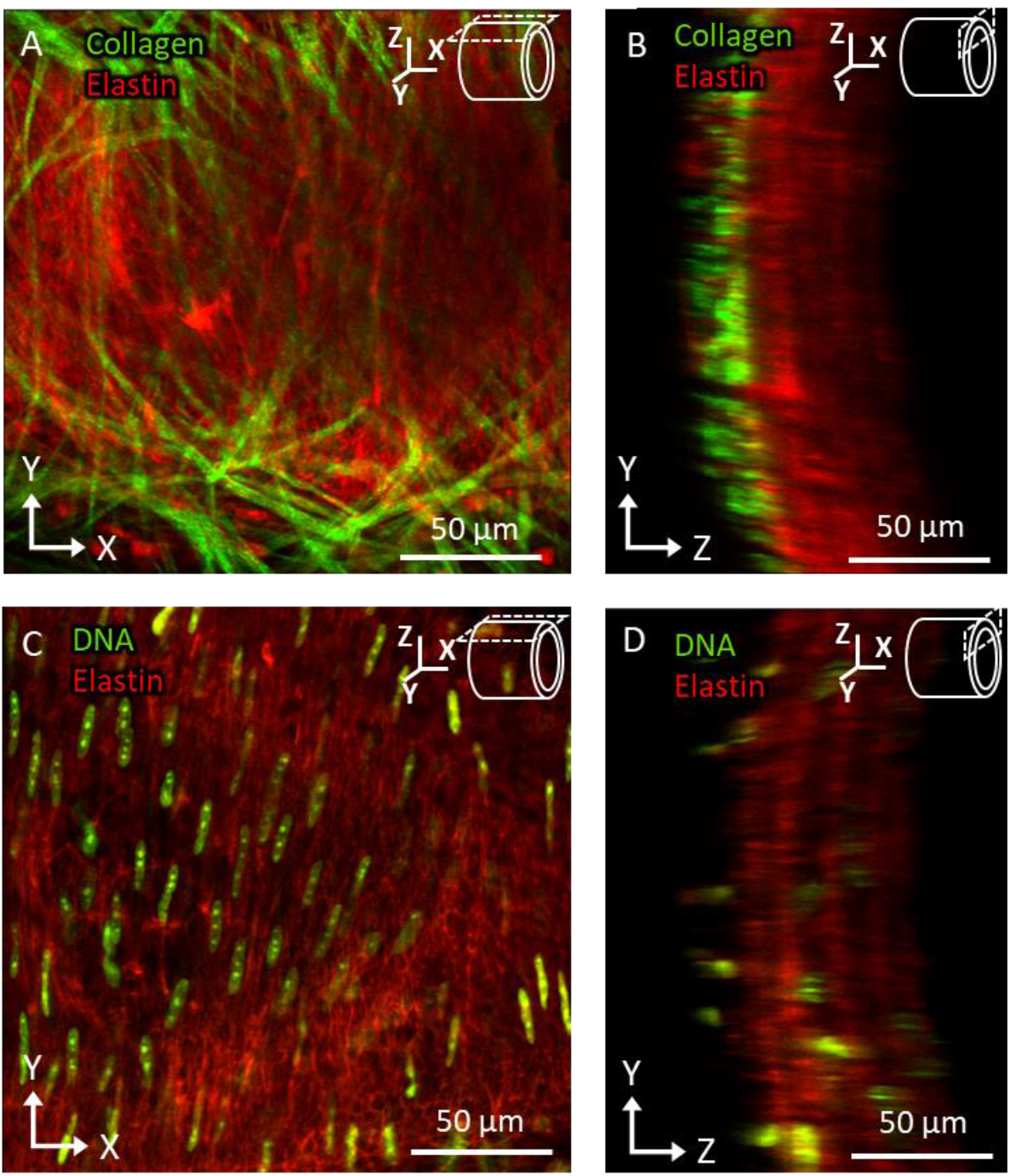
A, B: Collagen and elastin structure shown in green and red, respectively, with a top down and cross-sectional view with respect to the sample orientation. C, D: Nuclei and elastin structures shown in green and red, respectively, with a top down and cross-sectional view with respect to the sample orientation.

## Discussion

The described setup demonstrated its ability to replicate physiological loading conditions, enabling the examination of samples at their *in vivo*-like length and subjecting them to pulsatile pressure conditions with a peak-to-peak range of 40 mmHg and sinusoidal frequencies up to 20 Hz. Furthermore, the setup facilitated quasi-static and dynamic biomechanical characterization, vasoconstriction measurements, and capture of arterial wall constituent microstructure using TPLSM.

### Biomechanical characterization

Biomechanical characterization results captured the nonlinear biaxial elastic behaviour of the arterial wall, showing a stiffening effect as mechanical load reached and exceeded physiological ranges. It should be noted that measured axial force and axial stress appeared to have a strong dependence on both circumferential load and axial stretch, while circumferential stress, PWV, and inner diameter appeared to be dependent primarily on circumferential strain. In addition, the setup captured the significant viscoelastic stiffening effect when dynamically pressurizing arteries (28). While viscoelastic stiffening was significant for all the tested pulsatile pressurization frequencies compared to quasi static pressurization, stiffening between the tested frequencies was insignificant, indicating that much of the measured viscoelastic stiffening effect takes place at strain rates below the tested range. Together, these results demonstrate the importance of investigating arterial stiffening at well-defined axial stretches and under dynamic pressurization conditions that closely mimic physiological loading conditions, as deviation from these conditions can significantly alter arterial biomechanical behaviour. Furthermore, with this control over biaxial loading conditions, the presented setup can provide additional context for arterial stiffening research that deviated further from physiological loading conditions. It is this capacity to closely mimic physiological loading conditions that differentiates our setup from other platforms for investigating arterial stiffness (26, 29, 30).

### Advances over previously presented setup

Whilst sharing many features, the biaxial biomechanical characterization setup presented in this study has several key advantages over the previous biaxial biomechanical tester presented by our group (22). To start, the intraluminal and organ bath buffers are now separated, making it possible to add fluorescent dyes or vasoactive agents to the organ bath without contaminating the setup’s internals. In addition, now both the proximal and distal pipette positions are controlled using motorized stages, thus keeping the middle section of the sample within the field of view of the HSC or microscope when stretching the sample. Furthermore, the HSC provides improved axial and lateral resolution compared to the ultrasound transducer used previously and allows for diameter tracking during quasi-static stretch experiments. Lastly, the new pressurization scheme, consisting of a two-valve pressure regulator and custom pressure pulser, gives greater control and range for both quasi-static and dynamic pressurization of the sample. Together, these improvements enhance biomechanical characterization and provide new possibilities for investigating vasoreactivity and tissue microstructure.

### Study limitations

The long duration of the experiment protocol (∼5 hours) limits sample throughput, resulting in two or three samples being processed per day. In addition, because pressure containment was crucial for dynamic pressurization experiments, considerable time was spent on ensuring each sample was securely mounted in the setup without leaks.

Accurately and consistently determining unloaded wall thickness is important for calculating inner sample diameters from outer sample diameters. Consequently, cross-sectional rings for determining wall thickness needed to be cut as thin and perpendicular to the axis of the artery as possible. Failing to do so could result in overestimation of the wall thickness as more of the ring’s sidewall becomes visible in the cross-sectional image, which could be mistaken for the ring’s cross-sectional area. Furthermore, it is crucial to have the same individual conduct the image analysis process to determine wall thickness, as this helps mitigate the introduction of variation, given that it involves a partially manual procedure.

### Recommendations

Having demonstrated the capacity to investigate arterial vasoreactivity, biomechanical characteristics, and microstructure in one comprehensive setup with well-controlled loading conditions, our innovative setup opens new possibilities for interdisciplinary research into arterial remodelling and ageing-induced stiffening. Current evidence shows that VSMC contractility is dependent on axial stretch as well as dynamic circumferential strain (23, 25). With presented setup, the combined effect of axial stretch and dynamic pressurization on VSMC contractility can be investigated in depth, to further our understanding of how arterial wall loading impact on VSMC contractility, and vice versa.

In addition, the setup enables integrated characterization of constituent microstructure in relation to dynamic biomechanics, again under the same controlled conditions. For instance, in models of arterial aging, the interplay and contributions of elastin degradation and VSMC phenotypic modulation, as well as collagen growth and remodelling can now be studied in depth and linked to *in vivo*-like haemodynamics. Such investigations may prove ground-breaking in the field vascular aging and disease.

## Conclusion

The presented multi-modal characterization platform enables comprehensive *ex vivo* measurements under well-controlled *in vivo*-like loading conditions, for in-depth studies focusing on arterial stiffening. Our illustrative findings do emphasize the crucial need for controlling dynamic pressure and axial stretch conditions in investigating mechanistic and constitutive aspects of arterial stiffening.

